# Can early exposure to stress enhance resilience to ocean warming in two oyster species?

**DOI:** 10.1101/2020.01.21.913715

**Authors:** Roberta R. C. Pereira, Elliot Scanes, Mitchell Gibbs, Maria Byrne, Pauline M. Ross

## Abstract

Securing economically and ecologically significant molluscs, as our oceans warm and acidify due to climate change, is a global priority. South eastern Australia receives warm water in a strengthening East Australia Current and so resident species are vulnerable to elevated temperature and marine heat waves. This study tested whether oysters pre exposed to elevated temperature or heat stress enhances resilience to ocean warming later in life. Two Australian species, the flat oyster, *Ostrea angasi,* and the Sydney rock oyster, *Saccostrea glomerata*, were given a mild dose of warm water or “heat shock” stress in the laboratory and then transferred to elevated temperature conditions where we used the thermal outfall from power generation as a proxy to investigate the impacts of ocean warming. Shell growth, condition index, lipid content and profile and survival of oysters was impacted by elevated temperature in the field, with flat oysters being more impacted than Sydney rock oysters. Flat oysters grew faster than Sydney rock oysters at ambient temperature, but were more sensitive to elevated temperature. Early exposure to heat stress did little to ameliorate the negative effects of increased temperature, although the survival of heat shocked flat oysters was greater than non-heat shocked oysters. Further investigations are required to determine if early exposure to heat stress can act to inoculate oysters to future stress and overall enhance resilience of oysters to ocean warming.

## 1. Introduction

Climate change, the result of anthropogenic activities such as the burning of fossil fuels and deforestation, has exponentially increased the concentration of carbon dioxide (CO_2_) and other greenhouse gasses in the atmosphere [1]. Since the onset of the industrial revolution, atmospheric partial pressure of CO_2_ (*p*CO_2_) has increased from 280 ppm to 410 ppm causing global warming with direct impacts on the oceans [1, 2]. As a result, the world’s oceans have warmed by 0.68°C and for the East Australian coast are predicted to increase by up to 4°C by 2050 and 6°C before 2100 [3, 4]. Ocean warming and the increased incidence of heatwaves (abnormal high temperatures over multiple days [5]) negatively impacts diverse species [6]. Between 1925 and 2016 there has been a 54% annual increase in the duration of marine heatwaves worldwide [7]. Climate change is also impacting ocean stratification, currents, salinity, pH, sea level and increasing the frequency of extreme events [1,7,8].

Increasing frequency of thermal stress events will have consequences for fitness and survival of marine species and there is concern for habitat engineers such as bivalves and oysters [6, 9]. If oysters and other molluscs are to persist during this century along the southeast coast of Australia and in similar “hot spots” around the globe, they will need to be resilient to marine heat waves and habitat warming. It has been suggested that organisms can build resilience to environmental stress, through exposure to stress in early life. Studies have found that exposure to a mild stress early in life can result in later life stress resistance [10, 11]. Rather like a vaccination or inoculation, resistance to stress after exposure to mild stress in early life has been observed in a diverse array of organisms such as bacteria, plants, insects, mammals and fish [10, 11, 12, 13,14].

An increase in resistance to stress has been shown in the tidepool fish *Oligocottus maculosus* which after exposure to a +12°C heat stress had greater survival rates when exposed to subsequent stressful levels of high salinity and low oxygen concentration compared with fish that did not experience the heat shock [10]. The magnitude of the shock and recovery time played an important role in the stress response later in life [10]. Baltic Sea mussels *Mytilus edulis*, exposed to heat shock (+16°C) and then exposed to cadmium (20 µg L^-1^) produced heat shock proteins at a faster rate than mussels not exposed to heat stress [15]. Stress resistance may be enabled by production of protective heat shock proteins (e.g. HSP 70), although this is energetically costly. The mechanisms behind stress inoculation, are complex and likely not limited to production of heat shock proteins. Other processes such as alterations in metabolism and epigenetics are also thought to be involved [16, 17].

While mobile species can migrate changing their distribution as the ocean warms, sessile species are vulnerable because they are unable to move and the dispersive larval stages are often short-lived [18, 19]. It is predicted that sessile organisms such as oysters, which form the basis of aquaculture across the globe, will be impacted by elevated temperature, because of the energetic cost to physiological performance from climate change stress [20, 21]. Already, significant mortality has been reported for the north American oyster *Crassostrea virginica* exposed to elevated temperature, due to impacts on energetic reserves [22]. Reduced gametogenesis in *M. galloprovincialis* has been directly connected to warming [23]. Parental exposure to stress (in this case ocean acidification has, however, been shown to increase resilience of larval oysters, and this trait was carried over to adulthood [24, 25].

The flat oyster, *Ostrea angasi* and the Sydney rock oyster *Saccostrea glomerata* are native to south eastern Australia [26, 27], where they historically formed extensive reefs and are the basis of a USD $30 million aquaculture industry [28, 29]. *Saccostrea glomerata* is an intertidal species that occurs along the east and west coast of Australia with a current upper sea surface temperature (SST) range of 24-26 °C [28]. *Ostrea angasi* is distributed in shallow subtidal sheltered waterways along a similar range with a current upper SST temperature range of 22-24°C [27], however, this northern (warm) range is likely curtailed by historic overharvesting and introduced parasites in New South Wales (*Polydora* spp.)[27]. *O. angasi* are mostly found subtidally in comparatively stable thermal conditions [30]. These species are both currently the focus of reef restoration efforts along the south eastern coastline of Australia [31, 32] and are known to be vulnerable to acidification [33,34,35] and warming [33, 36]. South-eastern Australia receives warmer waters from the Coral Sea via the East Australian Current (EAC), which is strengthening [37, 38]. This region is considered a “hot spot”, as the rise in mean temperatures will be 3-4 times higher than the average for the world’s oceans and is also prone to marine heat waves [7,38,39].

The purpose of this study was to test the hypothesis that early exposure to heat stress or heat shock can be used as a mechanism to build resilience of *O. angasi* and *S. glomerata* to subsequent long-term exposure to warmed seawater. We used the thermal outfall from a power generating station as a proxy for ocean warming conditions as in previous studies [40]. Due to their different thermal ranges, distributions and habitats we predicted that *S. glomerata* will be more resilient than *O. angasi* to elevated temperature. As momentum gains to restore oyster reefs [31], knowledge of oyster responses and how to build resilience is needed to ensure sustainability of restoration efforts and the aquaculture industry.

## Methods

*Ostrea angasi* and *Saccostrea glomerata* were obtained from an oyster farm at Merimbula Lake (Merimbula Gourmet Oysters; 36°89’ 85”S, 149°88’ 46”E) and approximately 200 oysters per species were transported to Port Stephens Fisheries Institute (PSFI; 32°44’47”S, 152°03’30”E), following the protocol for oyster movement in New South Wales, Australia, during the Austral autumn 2018. The initial mean shell height was 69.68 ± S.E. 0.34 mm for *O. angasi* and 69.86 ± S.E. 0.33 mm for *S. glomerata*. After arrival at PSFI the oysters were placed in 40L tubs with seawater supplied from a 750L tank at 20°C. This temperature was the same as in Merimbula Lake when oysters were collected. Oysters were fed a mixture of microalgae cultured on-site containing 50 % *Chaetoceros muelleri* and 50 % *Tisochrysis lutea* at a concentration equivalent to 2 x 10^9^ cells oyster^-1^ d^-1^ [41] The initial mean (± S.E.) condition index for *O. angasi* and *S. glomerata* were 4.12 ± 0.42 g and 4.30 ± 0.39 g (n=6), respectively (see below for methods).

### 2.1 Heat shock

To determine if exposure to heat shock would confer subsequent resilience to long term exposure to elevated temperature, the following heat shock protocol was used. The oysters were divided into two sub-groups; one “control” and a “heat shocked” group per species into 750L tanks. Heat shock was administered by exposure to an elevated temperature of 26 °C for 18 hours and then 28°C for 6 more hours by slowly ramping up the temperature using aquarium heaters (Titan G2 1500 W). This was an initial +6°C (from 20° to 26°C) and a further increase of +2°C (from 26°C to 28°C). There was no mortality following heat shock treatment. Following the 24 hours at elevated temperature, the water was left to slowly cool to ambient (20°C). Oysters were submerged in ambient water overnight in the laboratory. On the following day, they were placed in baskets and left submerged at ambient conditions, in the adjacent estuary of PSFI (Tilligerry creek, Port Stephens) which remained at 20 °C for one week. After this period, they were removed and shell height was measured with a digital calliper.

A total of 40 oysters were randomly placed in baskets (600 x 250 x100 mm) divided into four compartments with 10 “control” *O. angasi* and 10 “control” *S. glomerata* which were exposed to ∼20°C at all times, 10 “heat shocked” *O. angasi* and 10 “heat shocked” *S. glomerata*, which were exposed to elevated temperature for 24 hours. The baskets were transported and deployed into Lake Macquarie (33°.07’94”, 151°.54’85”, Figure 1).

**Figure 1.**
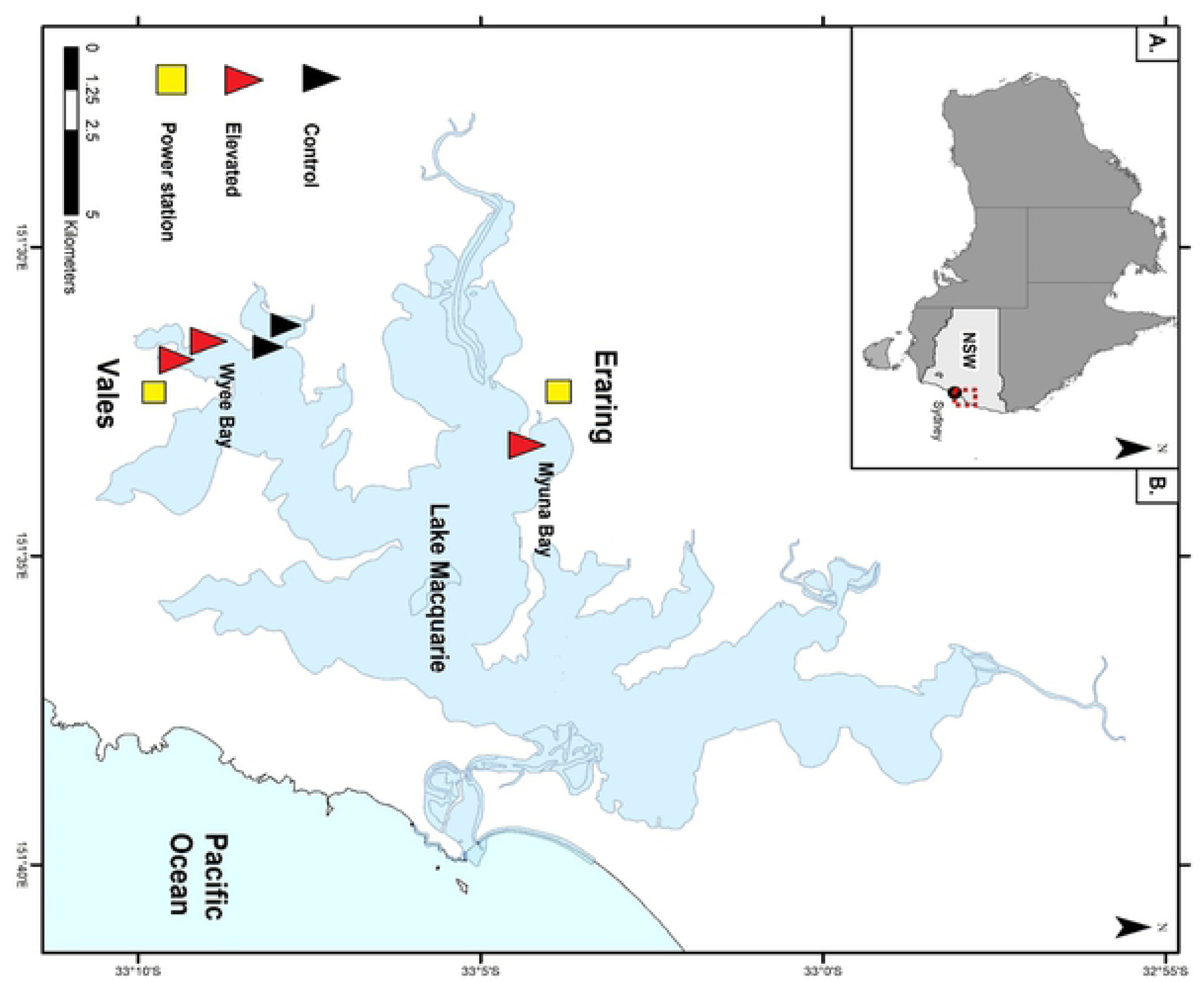
(A) Map of Australia with the study area in red (New South Wales, NSW). (B) Map of Lake Macquarie, NSW showing the field locations where the baskets were deployed for approximately seven months. Yellow squares represent the warm seawater outfall of two power stations (Eraring and Vales power stations). Black triangles are the ambient (control) locations and the red triangles are the elevated locations (total 5 baskets).

### 2.2 Field location

To determine the response of oysters in the real world of elevated temperature, we used warmed water released into a saline coastal lake by two power stations at Lake Macquarie, NSW. Lake Macquarie is a large coastal body of water in the centre of East Australian warming “hot spot”. Lake Macquarie is connected to the ocean and has daily tidal exchange. There is little freshwater input from the surrounding catchment [42]. Two coal fired power stations are located 23 kilometres apart on the shore of Lake Macquarie. Eraring power station is located in Myuna Bay (33° 4’2.92”S, 151°33’19.13”E) and Vales Point power station is located in Wyee Bay (33° 9’30.65”S, 151°31’48.37”E). Both stations use seawater from Lake Macquarie for cooling. The seawater is circulated for cooling and then released back into the estuary with no other treatment at a maximum of 37.5 °C as per licence requirements (NSW Environment Protection Licences 761; 1429). One location was selected near each power station outfall of the Eraring and Vales Point power stations in Lake Macquarie during May 2018 (autumn). A control location was also selected that represented the ambient mean temperature within Lake Macquarie which was not warmed by a power station. At each location, two baskets were deployed within 20m of each other. Each individual basket was attached to a 10 Kg concrete brick and contained a total of 40 oysters from both species and treatments (control/non heat shock and heat shock) and were deployed at a depth of 1.10 m by boat.

Temperature data were collected every 30 minutes by waterproof Hobo loggers (HOBO MX Pendant Temperature, Onset) attached to the baskets. Study locations were visited five times over seven months (late autumn to early summer) to download temperature data and renew the loggers. Oysters were deployed in Lake Macquarie for approximately seven months. At the end of the deployment (7 months) five baskets were retrieved; two from the ambient (control) location and three from elevated locations; two from Wyee Bay near Vales power station and one from Rocky Point near Earing power station (120 oysters from elevated temperature and 80 oysters from ambient temperature). Once retrieved, shell growth, condition index, standard metabolic rate and survival of oysters was measured. Total lipid and profile were measured in the laboratory.

### 2.3 Shell growth and condition index

To determine if exposure to heat shock confers subsequent resilience to long term exposure to elevated temperature on growth and condition index. measurements of oysters were done at the end of seven months of exposure in the field experiment.

There was no difference between shell height of oysters randomly allocated into heat shock and control (non-heat shock) treatments (One-way ANOVA comparing heat shock vs non-heat shock for each species, n=60; *O. angasi* = p > 0.05; *S. glomerata* = p > 0.05) at day zero. Final shell growth was then calculated as the difference between the final size of each individual oyster at seven months from an overall initial mean size of oysters per basket (n=10). The difference in shell growth was calculated by the formula:

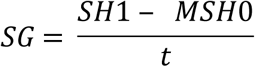

Where shell growth (SG) is the difference between final individual shell height (SH_1_) in millimetres and the mean initial shell height (MSH_0_) divided by time (t) in days.

The condition index of oysters was measured at the end of the experiment. Oysters were shucked, and body tissue and shell of individuals were dried in oven at 60°C for two days, to determine the dry weight (grams). The condition index (Ci) of oysters was then calculated by the formula [43, 44]:

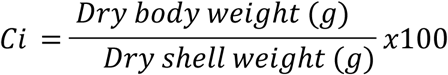

### 2.4 Standard Metabolic Rate (SMR)

To determine if exposure to heat shock would confer subsequent resilience to long term exposure to elevated temperature on standard metabolic rate (SMR), the SMR of 9-11 oysters of each species, treatment and basket (total 51 oysters; heat shock and control/non-heat shock) were measured at the end of the experiment using the methods of Parker et al. [33]. Measurements were done adjacent to the locations of collection to minimise stress of transport and to use seawater from Lake Macquarie.

To calculate SMR, oxygen consumption was measured by a closed respirometry system (OXY-10 PreSens, AS1 Ltd, Regensburg, Germany). Seawater was collected from Lake Macquarie and filtered through 0.47 µm glass filter paper before being used to fill respirometry chambers. Respirometers were built to accommodate the maximum oyster size (745ml and 830 ml). Each respirometer was connected to a fibre optic probe for measurement of dissolved oxygen in seawater. The probe was previously calibrated using two O_2_ concentration points (0% and 100% oxygen saturation of seawater) following the methods of Parker et al. [33]. Oysters were gently cleaned of any fouling organisms before placed in filtered seawater (adjusted to the corresponding treatment levels). The time that individuals took to lower the oxygen concentration in 20 % (∼1.2 O_2_ mg L^-1^) was recorded. Following the procedure of Parker et al. [33], only the time that the oyster is open and actively respiring (determined by observed decreasing oxygen) is used to calculate SMR. This is done to guard against the oyster remaining closed from handling stress. After each trial, each container was rinsed clean with filtered seawater (0.47 µm) and wiped clean with paper towel. After measurement the oysters were removed from the container and shucked to separate body tissues and shell. The tissue was then dried in an oven at 60°C for three days to measure their constant dry body tissue and shell weight in grams (±0.0001g, Analytical Balance Sartorius Research). Standard metabolic rates (SMR) were calculated by the formula:

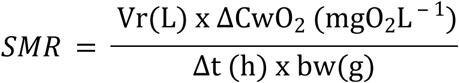

where SMR is the oxygen consumption normalized to 1 g of dry tissue mass (mg O_2_ g^-1^ dry tissue mass h-1, *V_r_* is the volume of the respirometry chamber minus the volume of the oyster (L), *ΔC_w_O_2_* is the change in water oxygen concentration measured (mg O_2_L^-1^), *Δt* is measuring time (h) and *b_w_* is the dry tissue mass (g) of the oyster.

### 2.5 Total lipid and lipid profile

To determine if exposure to heat shock and elevated temperature influences energy allocation, total lipid and lipid profiles were analysed. Body tissues of the oysters were placed in centrifuge tubes and frozen for analysis of total lipid content and lipid classes. The tissues were kept at −22°C for transport and then stored at −80°C until analysis. The tissues were then freeze dried (Alpha 1-4 LSCbasic, Martin Christ, Germany) and weighed in a microbalance (±0.0001g; Sartorius CPA225D). Lipids were extracted overnight using a modified Bligh & Dyer [45] one-phase methanol-chloroform-water extraction (2:1:0.8 v/v/v). The phases were separated by the addition of chloroform-water (final solvent ratio, 1:1:0.9 v/v/v methanol-chloroform-water). The total solvent extract (TSE) was concentrated using rotary evaporation at 40°C.

An aliquot of the TSE was analysed using an Iatroscan MK VI TH10 thin-layer chromatography-flame ionization detector (TLC-FID) analyser (Tokyo, Japan) to quantify individual lipid classes [46, 47]. Samples were applied in duplicate to silica gel SIII chromarods (5μm particle size) using 1 μl micropipettes. Chromorods were developed in a glass tank lined with pre-extracted filter paper. The primary solvent system used for the lipid separation was hexane-diethyl ether-formic acid (60:15:1.5), a mobile phase resolving non-polar compounds such as steryl ester (SE), triacylglycerol (TAG), free fatty acids (FFA), monoacylglycerol (MAG), Diacylglycerol (DAG). After development, the chromorods were oven dried and analysed immediately to minimize absorption of atmospheric contaminants. The FID was calibrated for each compound class (phosphatidylcholine (PL), cholesterol (Chol), cholesteryl palmitate (SE), palmitic acid (FFA), monopalmitin (MAG), dipalmitin (DAG), tripalmitin (TAG)). Peaks were quantified on an IBM compatible computer using DAPA Scientific software (Kalamunda, Western Australia, Australia). TLC-FID results are generally reproducible with a coefficient of variance of up to 3.46% of individual class abundances [48].

### 2.6 Survival

Oyster survival was determined after seven months deployment by emptying baskets one section at a time (to avoid mixing) and counting the total number of live oysters.

### 2.7 Data analysis

Statistical analyses were done using PRIMER v6+ software using either a three or two factor nested PERMANOVA (PRIMER v6+). This analysis was selected because it is robust to unbalanced designs [49].

For shell growth, condition index, SMR, and total lipids, data were analysed using a three factor PERMANOVA with “heat shock” as fixed factor with two levels (heat shock or control), “temperature” as fixed factor with two levels (ambient and elevated), and “basket” as random factor with two levels (basket 1 and basket 2) nested in temperature and heat shock. The analysis used 9999 permutations and only results with significance lower than 0.05 were considered as statistically different. The percentage survival at seven months was analysed using a two factor PERMANOVA with heat shock as fixed factor with two levels (heat shock or control) and temperature as fixed factor with two levels (ambient and elevated).

The composition of lipid profiles were fourth root transformed to limit the influence of large numbers [49] and analysed using a four factor multivariate PERMANOVA using the same model as above; with heat shock as fixed factor with two levels (Heat shock or control), temperature as fixed factor with two levels (control and elevated), and basket as random factor with two levels (basket 1 and basket 2) nested in temperature and heat shock. The analysis used 9999 permutations and only results with significance lower than 0.05 were considered as statistically different.

## 3. Results

### 3.1 Temperature

The average temperature over seven months at the ambient location was 20.06°C ± 3.85 (mean ± S.D.) and the average temperature at the elevated locations was 24.56°C ± 4.59 (Figure 2). The highest daily temperature experienced by oysters deployed at elevated temperature locations was 32.81 ± 0.39°C (mean ± S.D) in summer (December) and lowest daily average for the ambient location during the experiment was 14.92 °C ± 0.65 (mean ± S.D) in winter (August).

**Figure 2.**
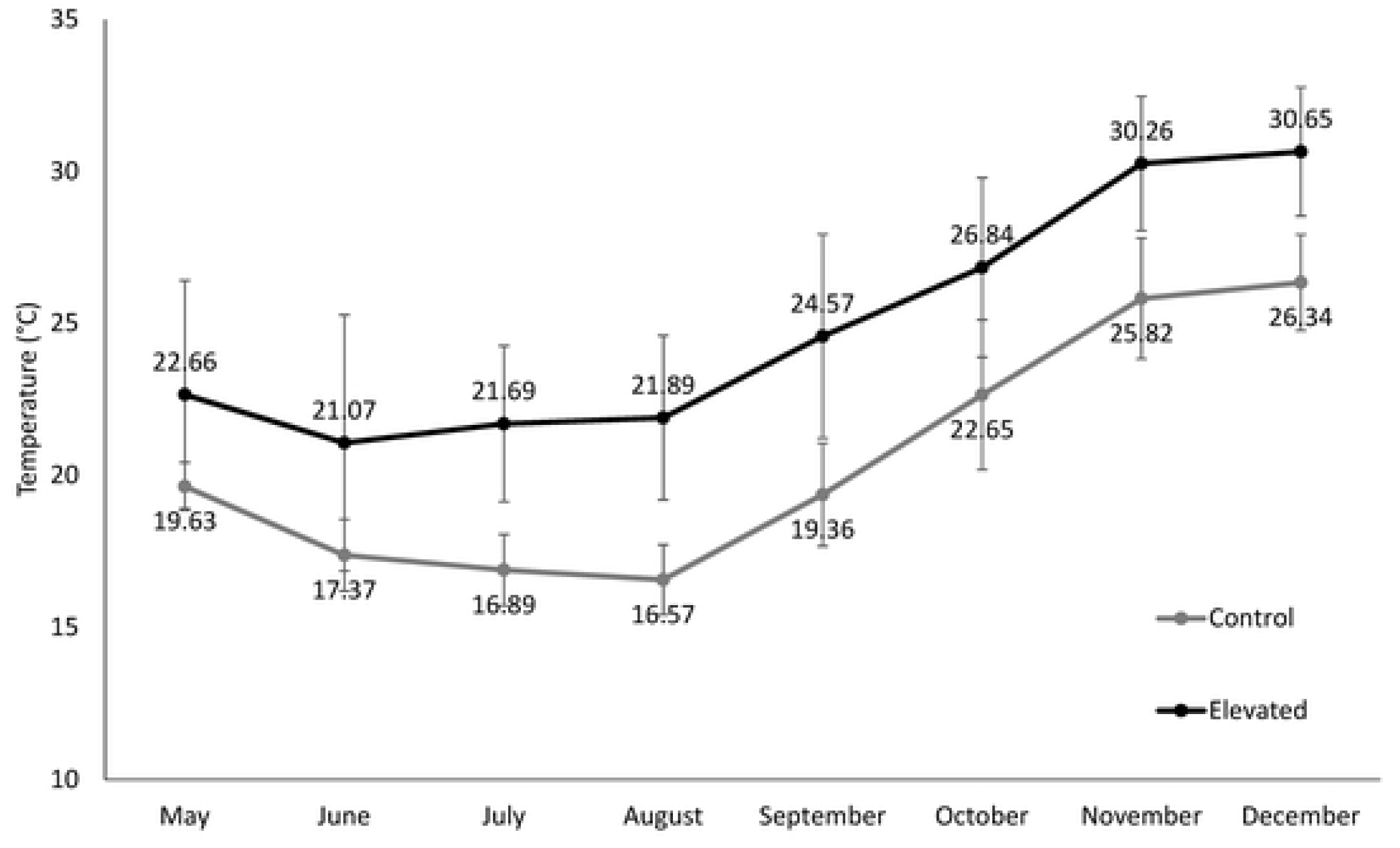
Mean monthly temperatures ± S.D. at control (ambient) and elevated temperature locations in Lake Macquarie, NSW from May to December 2018 (approximately seven months). Temperature data was measured every 30 minutes at 1.10m depth by water proof loggers.

### 3.2 Shell growth and condition index

Shell growth of *O. angasi* was almost ten-fold greater at ambient temperature compared to elevated temperature treatment (Figure 3a). Mean shell growth (mm day^-1^) was 0.10 ± 0.01 mm day^-1^ (mean ± S.E) at ambient temperature compared to 0.02 ± 0.01 mm day^-1^ and 0.01 ± 0.008 mm day^-1^ at elevated temperatures (Figure 3a). At ambient temperature, *O. angasi* which were heat shocked had lower growth than non-heat shocked oysters (Table 1a) and there was a trend for heat shocked *O. angasi* to have greater growth than non-heat shocked oysters at elevated temperature, but this was not significant. Shell growth of *S. glomerata* was not affected by temperature or heat shock (Figure 3b, Table 1b). *O. angasi* grew an order of magnitude greater than *S. glomerata* under ambient conditions, however, under elevated temperature there was little growth of either species (Figure 3 a, b).

**Figure 3.**
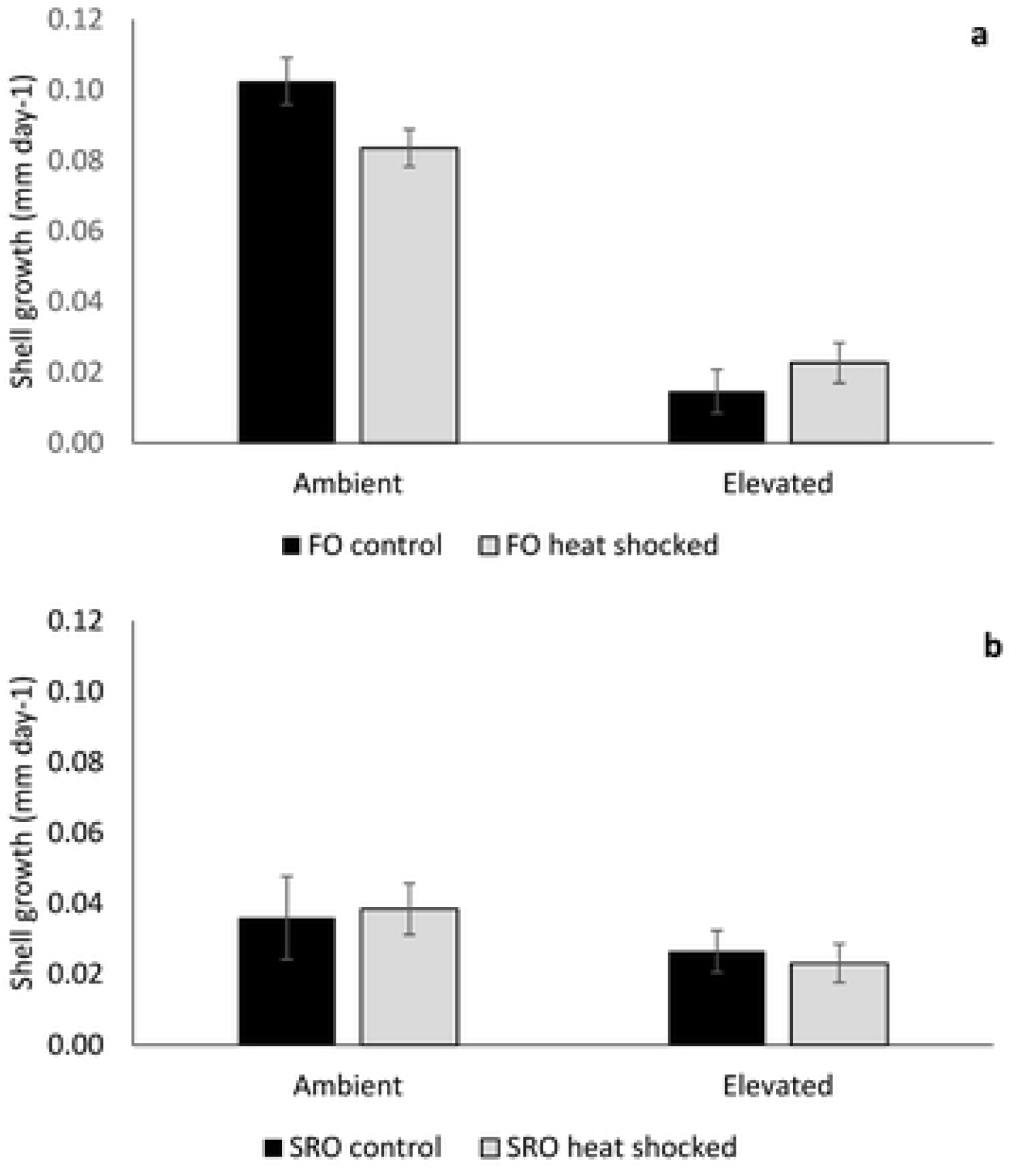
Mean difference in shell growth (± S.E.) for **a.** flat oysters, *Ostrea angasi* (FO control and FO heat shocked) and **b.** Sydney rock oysters, *Saccostrea glomerata* (SRO control and SRO heat shocked), exposed for seven months at ambient and elevated temperature locations at Lake Macquarie.

**Table 1a.** Shell growth, condition index and SMR of *Ostrea angasi* exposed for seven months in Lake Macquarie. P values were created using Monte Carlo tests.

|  | Shell Growth |  |  |  | Condition index |  |  |  | SMR |  |  |  |
| --- | --- | --- | --- | --- | --- | --- | --- | --- | --- | --- | --- | --- |
|  | df | MS | Pseudo-F | P(MC) | df | MS | Pseudo-F | P(MC) | df | MS | Pseudo-F | P(MC) |
| Heat Shock | 1 | 9.97 | 1.55 | 0.27 | 1 | 166.93 | 0.42 | 0.65 | 1 | 241.95 | 0.19 | 0.78 |
| Temperature | 1 | 3960 | 616.92 | **<0.001** | 1 | 3201.8 | 7.87 | **0.04** | 1 | 1748.70 | 1.36 | 0.3 |
| Heat Shock x Temperature | 1 | 165.88 | 25.84 | **<0.001** | 1 | 626.55 | 1.55 | 0.29 | 1 | 701.91 | 0.54 | 0.54 |
| Basket (Heat Shock x Temperature) | 4 | 5.91 | 0.2 | 0.94 | 3 | 402.87 | 1.56 | 0.20 | 4.00 | 1337.00 | 2.46 | 0.06 |
| Residuals | 69 | 29.23 |  |  | 28.00 | 258.09 |  |  | 20.00 | 543.10 |  |  |
| Total | 76 |  |  |  | 34.00 |  |  |  | 27.00 |  |  |  |

**Table 1b.** Shell growth, condition index and SMR of *Saccostrea glomerata* exposed for seven months in Lake Macquarie. P values were created using Monte Carlo tests.

|  | Shell Growth |  |  |  | Condition index |  |  |  | SMR |  |  |  |
| --- | --- | --- | --- | --- | --- | --- | --- | --- | --- | --- | --- | --- |
|  | df | MS | Pseudo-F | P(MC) | df | MS | Pseudo-F | P(MC) | df | MS | Pseudo-F | P(MC) |
| Heat Shock | 1 | 2.60 | 0.04 | 0.85 | 1 | 132.82 | 0.42 | 0.65 | 1 | 102.31 | 0.12 | 0.90 |
| Temperature | 1 | 186.07 | 2.84 | 0.17 | 1 | 2651.80 | 8.29 | **0.02** | 1 | 1181.50 | 1.39 | 0.29 |
| Heat Shock x Temperature | 1 | 0.00 | 0.00 | 0.99 | 1 | 777.29 | 2.43 | 0.16 | 1 | 1571.90 | 1.85 | 0.21 |
| Basket (Heat Shock x Temperature) | 4 | 66.49 | 1.49 | 0.22 | 4 | 319.59 | 0.96 | 0.46 | 4 | 880.30 | 1.38 | 0.24 |
| Residuals | 79 | 44.58 |  |  | 19 | 331.50 |  |  | 18 | 635.75 |  |  |
| Total | 86 |  |  |  | 26 |  |  |  | 25 |  |  |  |

The condition index of both *O. angasi* and *S. glomerata* was significantly lower at elevated temperature (Figure 4a,b, Table 1a,b) with no effect of heat shock treatment, although there was a slight trend for heat shocked *O. angasi* oysters at elevated temperature to have better condition.

**Figure 4.**
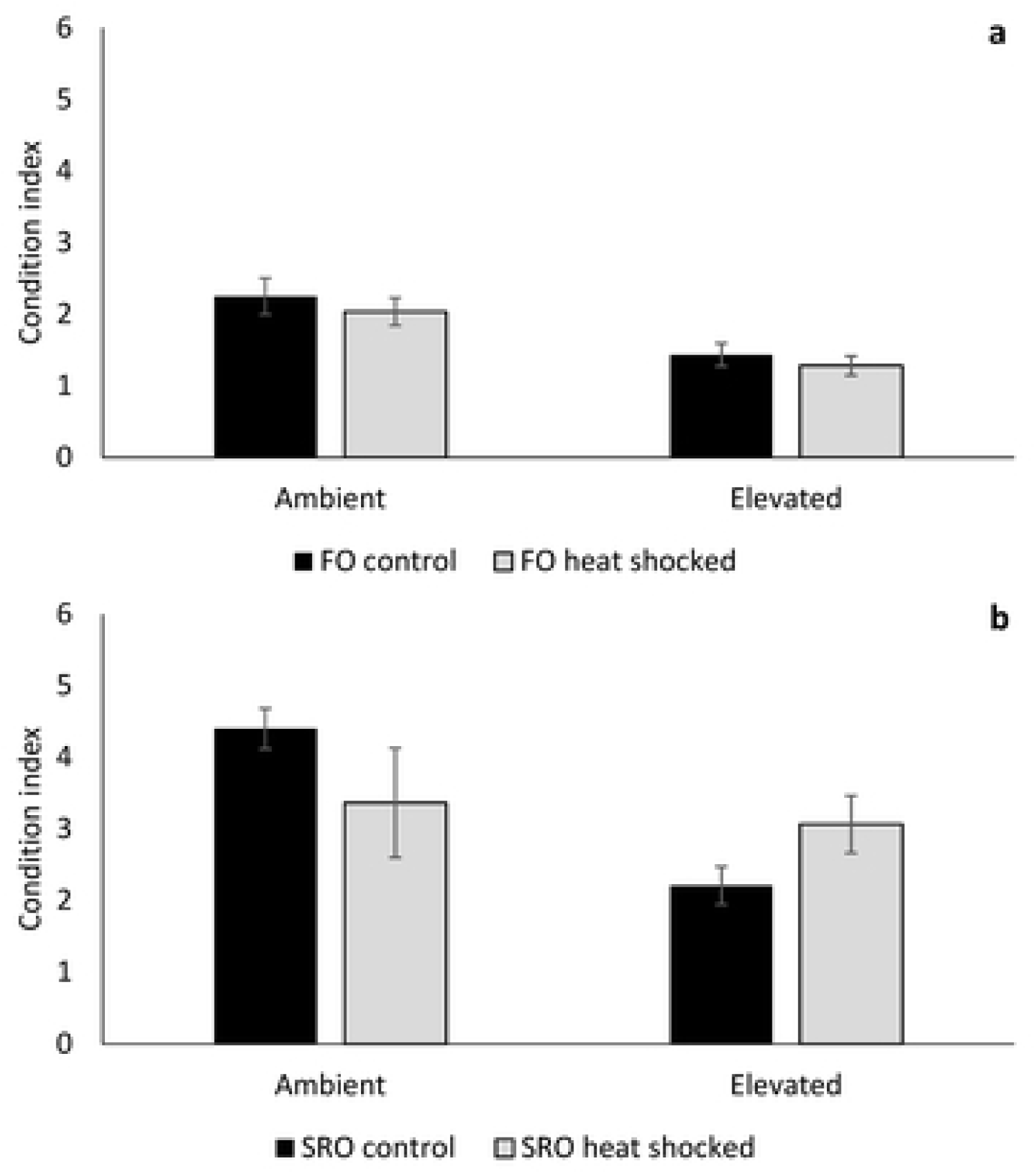
Mean condition index (± S.E.) of **a.** flat oysters, *Ostrea angasi* (FO control and FO heat shocked) and **b.** Sydney rock oysters, *Saccostrea glomerata* (SRO control and SRO heat shocked) exposed for seven months at ambient and elevated locations at Lake Macquarie.

### 3.3 Standard Metabolic Rate (SMR)

Standard Metabolic Rate of control, non-heat shocked *O. angasi* was lower at elevated temperature, but this was not significant (Figure 5a; Table 1a). SMR of control, non-heat shocked *S. glomerata* was greater at elevated temperature, but this was not significant (Figure 5b, Table 1b).

**Figure 5.**
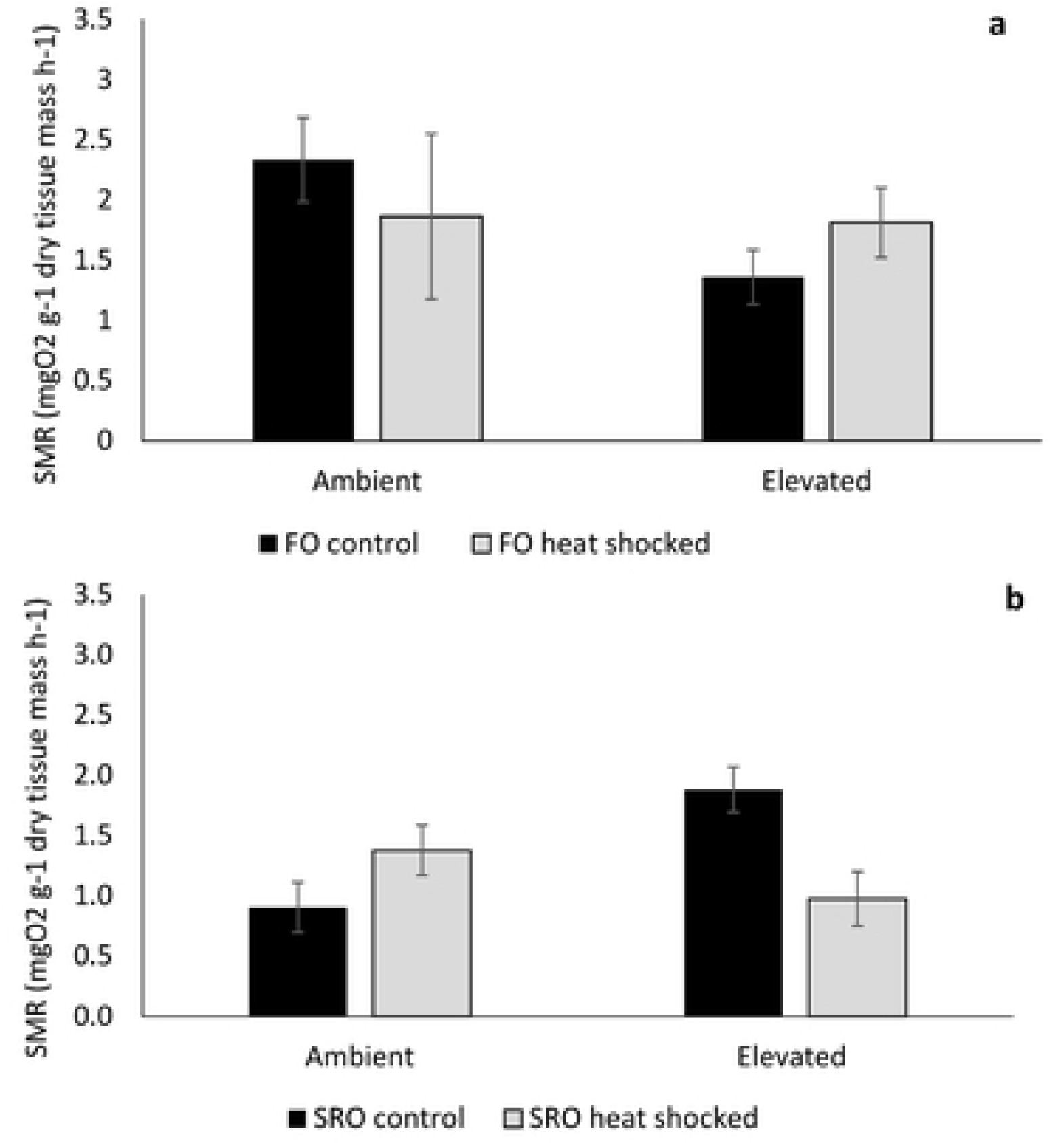
Mean standard metabolic rate (SMR) (± S.E.) of **a** flat oysters, *Ostrea angasi* (FO control and FO heat shocked) and **b.** Sydney rock oysters, *Saccostrea glomerata* (SRO control and SRO heat shocked) exposed for seven months at ambient and elevated temperature locations at Lake Macquarie.

### 3.4 Total lipids and lipid profiles

Mean total lipid content of *O. angasi* was greater at in those from the elevated temperature treatment compared to those held ambient (Figure 6a, Table 2a). There were no effects of heat shock or temperature on total lipid content in *S. glomerata* (Figure 6b, Table 2b). Lipid profile of *O. angasi*, was mostly driven by a greater amount of phospholipids in the oysters in the elevated temperature treatment (Figure 7a, Table 2a) and significantly lower amounts of TAGs (Figure 7a). Lipid profile of *S. glomerata* was similar across ambient and elevated temperatures, but there were significantly greater phospholipids at elevated temperature (Figure 7b, Table 2b).

**Figure 6.**
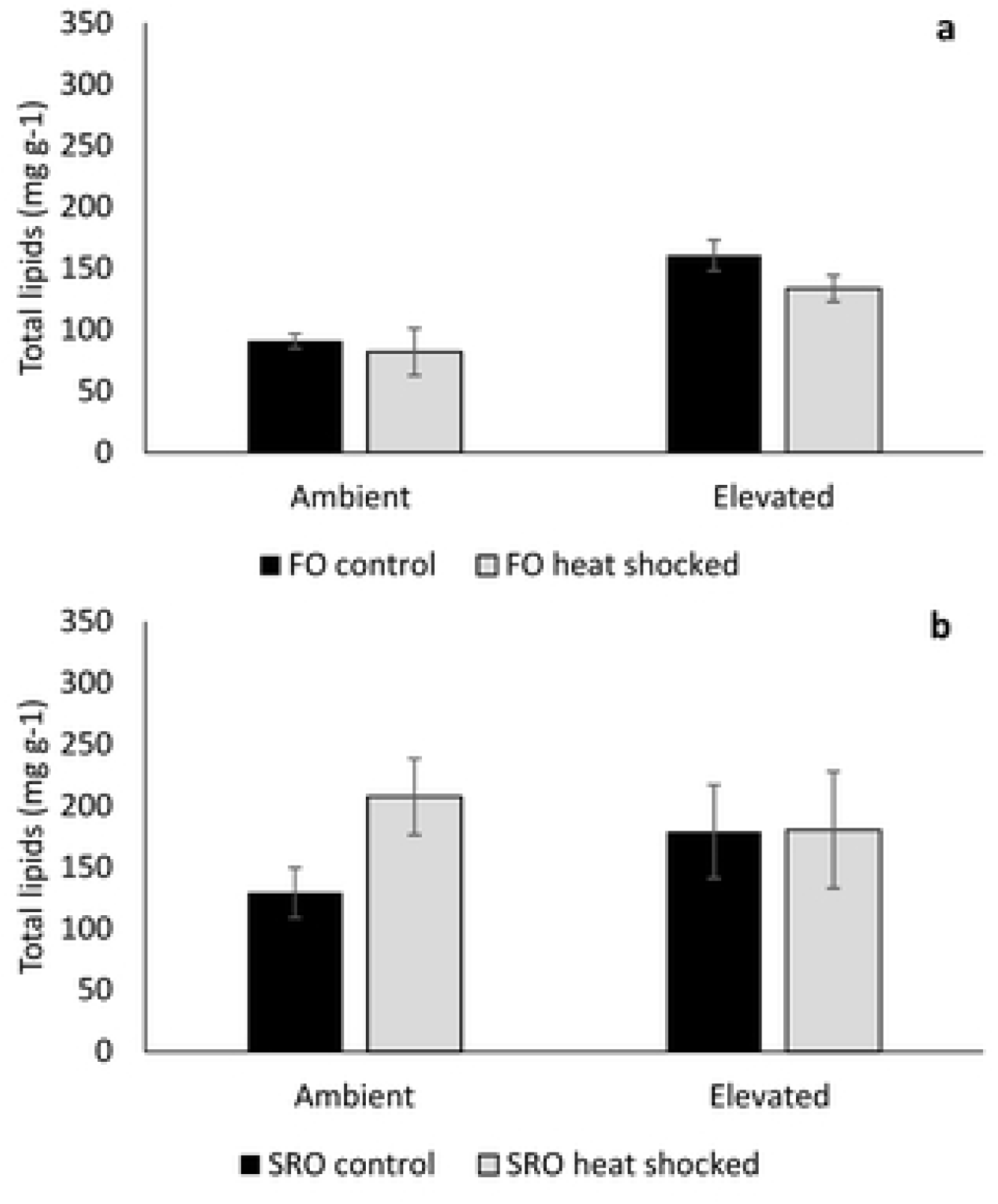
Mean total lipids (± S.E.) for **a.** flat oysters, *Ostrea angasi* (FO control and FO heat shocked) and **b.** Sydney rock oysters, *Saccostrea glomerata* (SRO control and SRO heat shocked) exposed for seven months at ambient and elevated temperature locations (n=5; except for HS oysters from Rocky Point – FO HS [n=3], SRO HS [n=2]) at Lake Macquarie.

**Figure 7.**
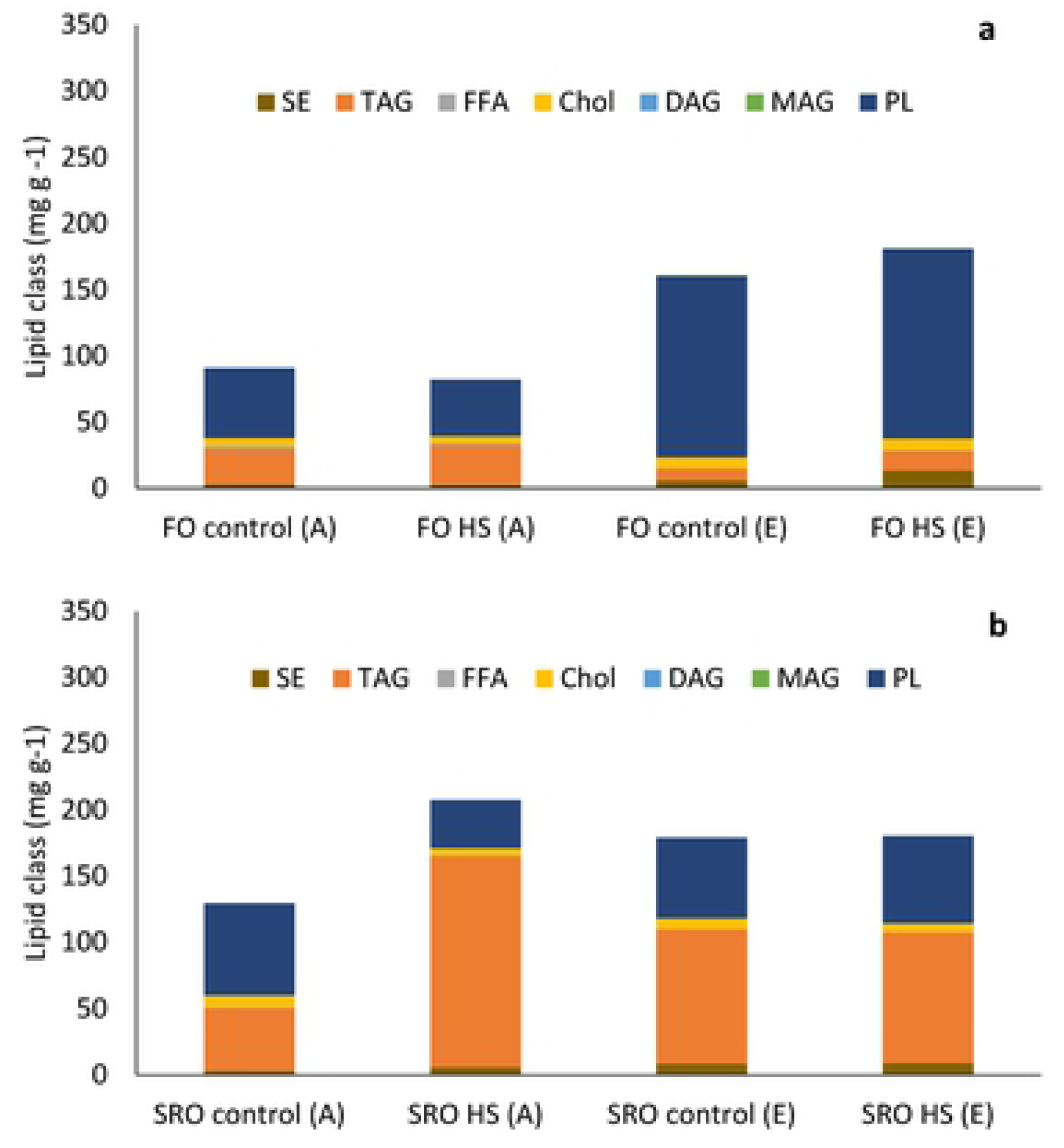
Lipid profile of **a.** flat oysters, *Ostrea angasi* (FO control FO heat shocked) after seven months exposure at ambient and elevated temperature locations and **b.** Lipid profile of Sydney rock oysters *Saccostrea glomerata* (SRO control and SRO heat shocked) exposed for seven months at ambient and elevated temperature. Lipid classes abbreviations are: SE – steryl ester; TAG-Triacyglyceride; FFA – Free Fatty Acids; Chol – Cholesterol; DAG – Diacylglyceride; MAG – Monoglyceride and PL – Polar Lipids.

**Table 2a.** Total lipids, amount of total lipids (mg/g), amount of Triacylglycerides (TAGs; mg/g) and Phospholipids (PLs; mg/g) of *Ostrea angasi* exposed for seven months in Lake Macquarie. P values were created using Monte Carlo tests.

|  | Total lipids |  |  |  | TAGs |  |  |  | PLs |  |  |  |
| --- | --- | --- | --- | --- | --- | --- | --- | --- | --- | --- | --- | --- |
|  | df | MS | Pseudo-F | P(MC) | df | MS | Pseudo-F | P(MC) | df | MS | Pseudo-F | P(MC) |
| Heat Shock | 1 | 204.36 | 0.58 | 0.55 | 1 | 3.58 | 0.02 | 0.96 | 1 | 780.33 | 1.20 | 0.36 |
| Temperature | 1 | 2652.8 | 7.55 | **0.03** | 1 | 798.48 | 3.65 | 0.13 | 1 | 8619.40 | 13.21 | **<0.001** |
| Heat Shock x Temperature | 1 | 270.63 | 0.77 | 0.48 | 1 | 72.98 | 0.33 | 0.62 | 1 | 347.69 | 0.53 | 0.64 |
| Basket (Heat Shock x Temperature) | 4 | 351.92 | 1.47 | 0.24 | 4 | 219.52 | 1.81 | 0.17 | 4 | 653.35 | 1.2 | 0.32 |
| Residuals | 14 | 239.88 |  |  | 15 | 121.39 |  |  | 15 | 545.05 |  |  |
| Total | 21 |  |  |  | 2 |  |  |  | 22 |  |  |  |

**Table 2b.** Total lipids (mg/g), amount of Triacylglycerides (TAGs; mg/g) and Phospholipids (PLs; mg/g), of *Saccostrea glomerata* exposed for seven months in Lake Macquarie. P values were created using Monte Carlo tests.

|  | Total lipids |  |  |  | TAGs |  |  |  | PLs |  |  |  |
| --- | --- | --- | --- | --- | --- | --- | --- | --- | --- | --- | --- | --- |
|  | df | MS | Pseudo-F | P(MC) | df | MS | Pseudo-F | P(MC) | df | MS | Pseudo-F | P(MC) |
| Heat Shock | 1 | 630.71 | 0.62 | 0.51 | 1 | 355.76 | 0.63 | 0.48 | 1 | 380.53 | 2.15 | 0.20 |
| Temperature | 1 | 585.15 | 0.57 | 0.54 | 1 | 40.98 | 0.07 | 0.86 | 1 | 1291.40 | 7.31 | **0.04** |
| Heat Shock x Temperature | 1 | 824.63 | 0.81 | 0.44 | 1 | 951.68 | 1.70 | 0.26 | 1 | 957.97 | 5.42 | 0.06 |
| Basket (Heat Shock x Temperature) | 4 | 1023.70 | 1.50 | 0.22 | 4 | 564.66 | 1.68 | 0.20 | 4 | 172.28 | 0.37 | 0.84 |
| Residuals | 17 | 682.38 |  |  | 17 | 337.06 |  |  | 17 | 461.41 |  |  |
| Total | 24 |  |  |  | 24 |  |  |  | 24 |  |  |  |

### 3.5 Survival

Survival of heat-shocked *O. angasi* was significantly greater than non-heat shocked oysters at ambient and elevated temperature (Figure 8a; Table 3a). At the ambient and elevated temperature locations, survival of *O. angasi* was greatest for the heat shocked oysters (Ambient, control oysters = 90% and heat shocked oysters= 100%; Elevated, control = 53% and heat shocked = 80%). Survival of *S. glomerata* was significantly lower for control oysters at ambient temperature compared to heat-shocked oysters (Figure 8b, Table 3b).

**Figure 8.**
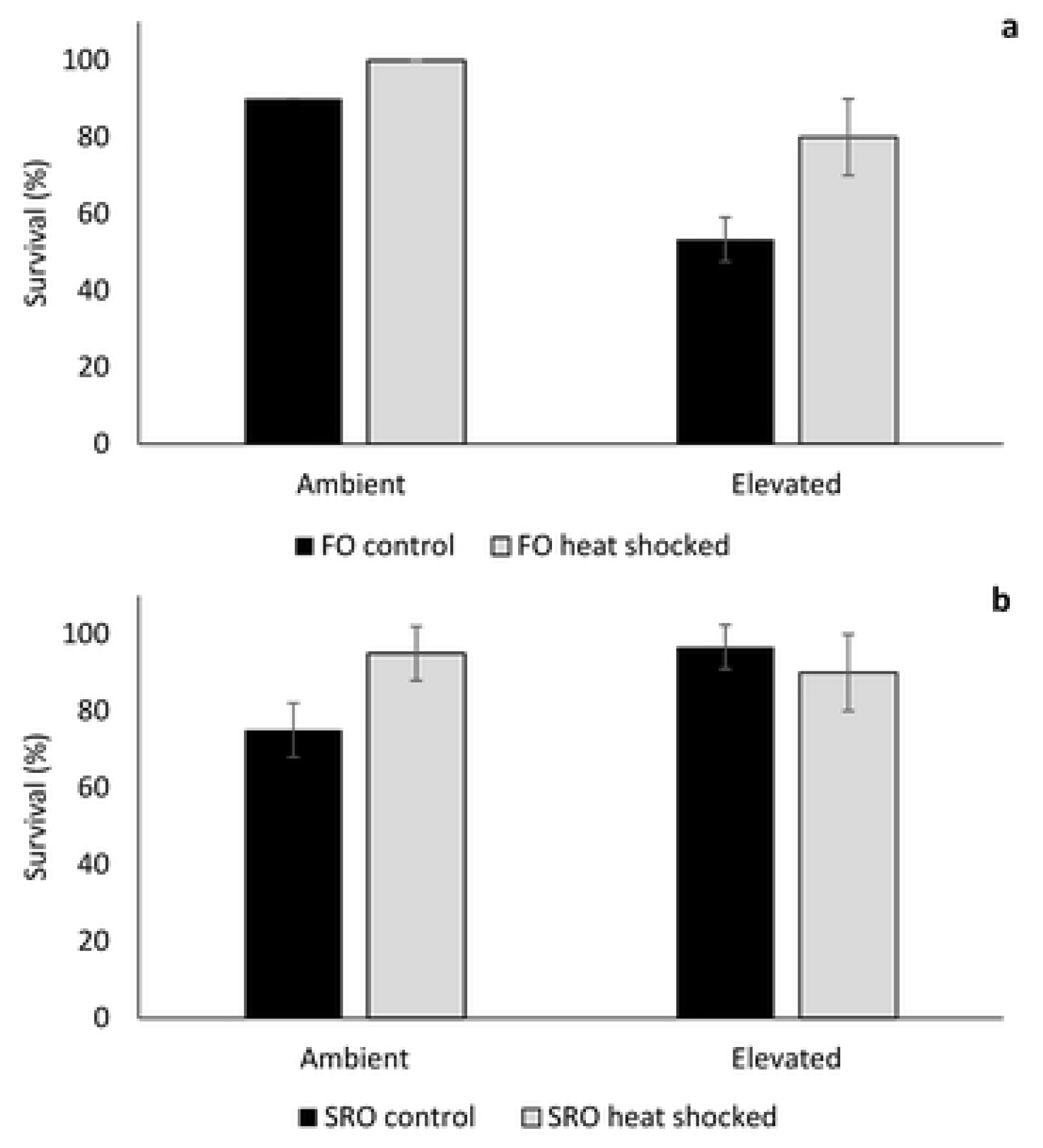
Mean survival (±S.D.) of **a** flat oysters, *Ostrea angasi* (FO control and FO heat shocked) and **b.** Sydney rock oysters, *Saccostrea glomerata*, (SRO control and SRO heat shocked), exposed for seven months at ambient and elevated temperature locations at Lake Macquarie.

**Table 3a.** Percentage survival of *Ostrea angasi* exposed for seven months in Lake Macquarie. P values were created using Monte Carlo tests.

|  | df | MS | Pseudo-F | P(MC) |
| --- | --- | --- | --- | --- |
| Heat Shock | 1 | 359.59 | 16.16 | **0.007** |
| Temperature | 1 | 802.23 | 36.05 | **0.001** |
| Heat Shock x Temperature | 1 | 142.45 | 6.40 | **0.04** |
| Residuals | 6 | 22.25 |  |  |
| Total | 9 |  |  |  |

**Table 3b.** Percentage survival of *Saccostrea glomerate* exposed for seven months in Lake Macquarie. P values were created using Monte Carlo tests.

|  | df | MS | Pseudo-F | P(MC) |
| --- | --- | --- | --- | --- |
| Heat Shock | 1 | 39.189 | 2.019 | 0.2037 |
| Temperature | 1 | 57.668 | 2.971 | 0.1309 |
| Heat Shock x Temperature | 1 | 143.35 | 7.3852 | **0.0353** |
| Residuals | 6 | 19.41 |  |  |
| Total | 9 |  |  |  |

## 4. Discussion

Exposure to long term warming in the field had negative impacts on shell growth, condition index, and survival of *O. angasi* and *S. glomerata*. Shell growth, condition index, lipid content and profile and survival, but not SMR of oysters was impacted by elevated temperature, with flat oysters more impacted than Sydney rock oysters. Flat oysters grew faster than Sydney rock oysters at ambient temperature, but were more sensitive to elevated temperature. Exposure early in life to heat shock did little to ameliorate the negative effects of elevated temperature, although there was a trend for shell growth and condition index and a significant effect of survival of heat shocked flat oysters to be greater than control oysters at elevated temperature. SMR was not significantly impacted by elevated temperature, although once again there was a trend for SMR of flat oysters to decrease with increased temperature and for SMR of Sydney rock oysters to increase with increased temperature. The lipid profile of *O. angasi* was also reduced by elevated temperature, while the lipid profile for *S. glomerata* was not affected.

As oysters are ectothermic organisms, changes in external temperature away from their optimum causes physiological processes to become less efficient and homeostasis begins to require more energy [50]. The effects of elevated temperature on *O. angasi* and *S. glomerata* are similar to those observed for other bivalve species. For example, Hiebenthal et al. [51] found lower growth and condition for *Arctica islandica* at elevated temperature (16°C) compared to the control (7.5°C) and an intermediate treatment (10°C). Condition index and survivorship of *M. edulis* was reduced under elevated temperature (25°C) compared with control [51]. Effects on these physiological processes, were attributed to thermal sensitivity of *A. islandica* to temperatures outside its distribution and to accumulation of lipofuscin, a disease related pigment [51]. For *Mercenaria mercenaria* and *Argopecten irradians*, elevated temperature (28°C) impacted shell growth of juveniles [52].

Elevations in temperature increase the SMR of marine ectotherms until a point known as the “Arrhenius Breakpoint Temperature” (ABT). When ABT is reached, SMR rapidly declines indicating that the organism can no longer meet their energetic requirements at that temperature [33]. The reduced growth of *O. angasi* at elevated compared to ambient temperature was correlated with a trend for lower SMR, indicating that *O. angasi* may have experienced temperatures beyond their ABT. Temperature had no effect on the SMR of *S. glomerata.* Parker et al., [33] found that increases in seawater temperature can increase the SMR of *S. glomerata.* SMR of *S. glomerata* increased with increased temperature up to 33 °C (the upper temperature treatment in that study) indicating an ABT for *S. glomerata* of above 33 °C. Increased SMR can impact energy budget and may indicate a thermal response with extra costs needed to cover basal metabolism [33, 53]. For oysters, thermal stress can also alter cardiac function, protein synthesis [53] and gametogenesis [23].

Oysters have the capacity to store surplus energy ingested from food in the form of lipids which can assist in the persistence during stressful conditions. While the lipid profile of *S. glomerata* was not impacted by elevated temperature, there were significant impacts of elevated temperature on total lipids and lipid profile, especially Triacylglycerides (TAGs) of *O. angasi.* TAGs are the primary source of stored lipid energy for bivalves [54], indicating that *O. angasi* had begun to use stored lipid reserves. Studies have found that under stressful conditions bivalves have lower lipid reserves. For example, exposure to elevated *p*CO_2_ decreased the lipid index of larvae of *A. irradians*, *M. mercenaria and C. virginica* which further declined when combined with warming [52, 55]. Lipid levels in the eggs of *S. glomerata* also decreased when exposed to the dual stress of elevated *p*CO_2_ and copper [56]. Further research on how lipids are used by oysters in response to stress could provide insights into the ramifications of living in warmer oceans.

### Stress inoculation, and resilience

This study tested the hypothesis that pre exposure of oysters to heat shock stress will build resilience to later exposure to elevated temperature. Stress inoculation leading to stress resilience has been observed in diverse phyla from bacteria to mammals e.g. [10, 11]. Heat shock may help to build resilience, but at the same time have costs. For example, heat shocked *O. angasi* had significantly greater rates of survival at elevated temperatures, but heat shocked oysters had less growth at ambient temperature. Perhaps energy was used to produce heat shock proteins or other protective measures, thereby reducing energetic reserves for growth and other important physiological processes.

Heat shocked *O. angasi* had greater rates of survival at elevated temperature, and a similar trend was observed for *S. glomerata* which had greater than 90% survival at elevated temperature. When organisms experience stressful temperatures, they undergo a thermal response, which is energy dependent [57]. This thermal response includes producing chaperones, such as energetically expensive heat shock proteins (HSPs) [57, 58]. Species with lower thermal tolerance might be induced to produce HSPs in response to elevated temperatures before more tolerant species, which can endure longer periods under warming stress (e.g. *M. trossulus and M. galloprovincialis;* [58]). Production of these molecular chaperones (HSPs) is a common response to elevated temperature [57]. Heat shock proteins have important functions when an organism is exposed to elevated temperature, including degradation of denatured proteins and prevention of misfolding, having a key function on cellular protection [59]. These responses (e.g. expression of heat shock proteins, antioxidants, increased respiration rates) all incur an energetic cost which can cause an imbalance in the energetic partitioning of individuals [33,57,60,61].

Overall, *S. glomerata* was found to be generally more tolerant of habitat warming than *O. angasi*. *S. glomerata* had no change in shell growth although they were in poorer condition at elevated temperature of 28-30 °C. As an intertidal species that experiences a highly dynamic thermal environment, *S. glomerata* could be expected to be more thermally tolerant as has been shown for other intertidal organisms [62, 63]. These findings are supported by previous work by Parker et al., [33] that showed 33 °C was not beyond their ABT, and the distribution of *S. glomerata* which extends further north along the east coast of Australia than *O. angasi*. Additionally, *S. glomerata* can experience air temperatures in excess of 40 °C during emersion at low tide [64]. The lack of effect of elevated temperature on *S. glomerata* indicates that they were not placed beyond their thermal limits in the deployment used here, in contrast to *O. angasi*, which did not cope as well.

*O. angasi* had the greatest growth rate at ambient conditions. The shell growth of *O. angasi* was over ten-fold greater than Sydney rock oysters after seven months, as expected from growth in aquaculture [65]. While *O. angasi* grew well at ambient conditions, growth and survival were impacted by warming. As this species lives in a relatively stable, sub-tidal habitat we expected this species to be more sensitive to warming compared to *S. glomerata*.

Globally and across Australia efforts are being made to restore oyster reefs [31,66,67]. Climate change will impact on oyster reef restoration [36]. Projected ocean warming for the region (4°C) as well as contemporary marine heat waves, as seen in the region recently [39] are an important consideration for reef restoration efforts along the south-eastern coastline of Australia. Our study has shown that using thermal outfall as a proxy for ocean warming can be useful for predicting future warming. This approach is similar to natural laboratories using underwater CO_2_ vents which have successfully tested the responses of marine organisms to ocean acidification [68, 69]. Our results indicate that habitat warming will be a greater threat to *O. angasi* compared to *S. glomerata.* As ocean warming will not act alone, oyster reef restoration is at risk from multiple stressors including ocean acidification, salinity, and other environmental pollutants which will act simultaneously [36]. These co-occurring stressors further threaten native species of oysters, other molluscs and marine organisms and so mitigation strategies to build oyster resilience will be critical. Our results indicate that early exposure to stress inoculation does not enhance resilience and may not be useful strategy, especially for restoration ventures involving *O. angasi*.

## Acknowledgements

We acknowledge and thank CNPq – Conselho Nacional de Desenvolvimento Científico e Tecnológico – Brazil (PhD CNPq scholarship) for financially supporting this work. We also would like to thank Port Stephens Fisheries Institute, Dr Wayne O’Connor and the Office of Environment and Heritage for great support during development of this study. We thank Kyle Johnston and Richard Grainger for assistance in the field.

## References

1. IPCC. Summary for policymakers. In: Climate Change 2014: Impacts, Adaptation, and Vulnerability. Part A: Global and Sectoral Aspects. Contribution of Working Group II to the Fifth Assessment Report of the Intergovernmental Panel on Climate Change [Field, C.B., V.R. Barros, D.J. Dokken, K.J. Mach, M.D. Mastrandrea, T.E. Bilir, M. Chatterjee, K.L. Ebi, Y.O. Estrada, R.C. Genova, B. Girma, E.S. Kissel, A.N. Levy, S. MacCracken, P.R. Mastrandrea, and L.L. White (eds.)]. Cambridge University Press, Cambridge, United Kingdom and New York, NY, USA. 2014:pp. 1–32.

2. Cheng L, Abraham J, Hausfather Z, Trenberth KE. How fast are the oceans warming? Science. 2019;363(6423):128–9.

3. Hobday AJ, Okey TA, Poloczanska ES, Kunz TJ, Richardson AJ. Impacts of climate change on Australian marine life. Report to the Australian Greenhouse Office, Canberra, Australia. 2006.

4. Lenton A, McInnes KL, O’Grady JG. Marine projections of warming and ocean acidification in the Australasian region. Australian Meteorological and Oceanographic Journal. 2015;65(1):1–28.

5. Hobday AJ, Alexander LV, Perkins SE, Smale DA, Straub SC, Oliver EC, et al. A hierarchical approach to defining marine heatwaves. Progress in Oceanography. 2016;141:227–38.

6. Ridgway K, Hill K. The East Australian Current. A marine climate change impacts and adaptation report card for Australia. 2009;5(09).

7. Oliver EC, Donat MG, Burrows MT, Moore PJ, Smale DA, Alexander LV, et al. Longer and more frequent marine heatwaves over the past century. Nat Commun. 2018;9(1):1324.

8. Doney SC, Ruckelshaus M, Duffy JE, Barry JP, Chan F, English CA, et al. Climate change impacts on marine ecosystems. Annu Rev Mar Sci. 2012;4:11–37.

9. Oliver EC, Benthuysen JA, Bindoff NL, Hobday AJ, Holbrook NJ, Mundy CN, et al. The unprecedented 2015/16 Tasman Sea marine heatwave. Nat Commun. 2017;8:16101.

10. Todgham AE, Schulte PM, Iwama GK. Cross-tolerance in the tidepool sculpin: the role of heat shock proteins. Physiological and Biochemical Zoology. 2005;78(2):133–44.

11. Parker KJ, Buckmaster CL, Sundlass K, Schatzberg AF, Lyons DM. Maternal mediation, stress inoculation, and the development of neuroendocrine stress resistance in primates. Proceedings of the National Academy of Sciences. 2006;103(8):3000–5.

12. Laplace JM, Boutibonnes P, Auffray Y. Unusual resistance and acquired tolerance to cadmium chloride in *Enterococcus faecalis*. J Basic Microbiol. 1996;36(5):311–7.

13. Krebs RA, Feder ME. Hsp70 and larval thermotolerance in *Drosophila melanogaster*: how much is enough and when is more too much? J Insect Physiol. 1998;44(11):1091–101.

14. Munne-Bosch S, Alegre L. Cross-stress tolerance and stress“memory” in plants. Environ Exp Bot. 2013;94:1–88.

15. Tedengren M, Olsson B, Reimer O, Brown DC, Bradley BP. Heat pretreatment increases cadmium resistance and HSP 70 levels in Baltic Sea mussels. Aquat Toxicol. 2000;48(1):1–12.

16. Chapple JP, Smerdon GR, Berry R, Hawkins AJ. Seasonal changes in stress-70 protein levels reflect thermal tolerance in the marine bivalve *Mytilus edulis* L. J Exp Mar Biol Ecol. 1998;229(1):53–68.

17. Huey RB, Bennett AF. Physiological adjustments to fluctuating thermal environments: an ecological and evolutionary perspective. In: Morimoto RI, Tissieres A, Georgopoulos C, editors. Stress proteins in biology and medicine. Cold Spring Harbor, NY: Cold Spring Harbor Lab. Press; 1990;19:37–59.

18. Pechenik JA. On the advantages and disadvantages of larval stages in benthic marine invertebrate life cycles. Mar Ecol Prog Ser. 1999;177:269–97.

19. Gattuso J-P, Magnan A, Billé R, Cheung WW, Howes EL, Joos F, et al. Contrasting futures for ocean and society from different anthropogenic CO_2_ emissions scenarios. Science. 2015;349(6243):aac4722.

20. Pörtner H-O. Integrating climate-related stressor effects on marine organisms: unifying principles linking molecule to ecosystem-level changes. Mar Ecol Prog Ser. 2012;470:273–90.

21. Sokolova IM. Energy-limited tolerance to stress as a conceptual framework to integrate the effects of multiple stressors. Integ and Comp Biol. 2013;53(4):597–608.

22. Ivanina AV, Dickinson GH, Matoo OB, Bagwe R, Dickinson A, Beniash E, et al. Interactive effects of elevated temperature and CO_2_ levels on energy metabolism and biomineralization of marine bivalves *Crassostrea virginica* and *Mercenaria mercenaria*. Comp Biochem Physiol A Mol Integr Physiol. 2013;166(1):101–11. doi: https://doi.org/10.1016/j.cbpa.2013.05.016.

23. Fearman J, Moltschaniwskyj N. Warmer temperatures reduce rates of gametogenesis in temperate mussels, *Mytilus galloprovincialis*. Aquaculture. 2010;305(1-4):20–5.

24. Parker LM, Ross PM, O’Connor WA, Borysko L, Raftos DA, Pörtner HO. Adult exposure influences offspring response to ocean acidification in oysters. Glob Change Biol. 2012;18(1):82–92.

25. Parker LM, O’Connor WA, Raftos DA, Pörtner H-O, Ross PM. Persistence of positive carryover effects in the oyster, *Saccostrea glomerata*, following transgenerational exposure to ocean acidification. PloS one. 2015;10(7):e0132276.

26. Scanes E, Johnston EL, Cole VJ, O’Connor WA, Parker LM, Ross PM. Quantifying abundance and distribution of native and invasive oysters in an urbanised estuary. Aquatic Invasions. 2016;11(4):425–36.

27. Ogburn DM, White I, Mcphee DP. The disappearance of oyster reefs from eastern Australian estuaries—impact of colonial settlement or mudworm invasion? Coast Manage. 2007;35(2-3):271–87.

28. Nell JA. The history of oyster farming in Australia. Marine Fisheries Review. 2001;63(3):14–25.

29. NSW Department of Primary Industries, Aquaculture Production Report 2010-2011. Port Stephens, New South Wales: Department of Primary Industries, 2011. ISSN 1444-840.

30. Crawford, C, National review of Ostrea angasi aquaculture: historical culture, current methods and future priorities, University of Tasmania Institute for Marine and Antarctic Studies, Hobart, Tasmania 2016.

31. Gillies CL, Crawford C, Hancock B. Restoring Angasi oyster reefs: What is the endpoint ecosystem we are aiming for and how do we get there? Ecol Manage Restor. 2017;18(3):214–22.

32. McLeod I, Boström-Einarsson L, Creighton C, D’Anastasi B, Diggles B, Dwyer P, et al. Habitat value of Sydney rock oyster (*Saccostrea glomerata*) reefs on soft sediments. Mar and Freshw Res.2019. https://doi.org/10.1071/MF18197

33. Parker LM, Scanes E, O’Connor WA, Coleman RA, Byrne M, Pörtner H-O, et al. Ocean acidification narrows the acute thermal and salinity tolerance of the Sydney rock oyster *Saccostrea glomerata*. Mar Pollut Bull. 2017;122(1-2):263–71.

34. Parker LM, Ross PM, O’Connor WA. Comparing the effect of elevated *p*CO_2_ and temperature on the fertilization and early development of two species of oysters. Mar Biol. 2010;157(11):2435–52.

35. Cole VJ, Parker LM, O’Connor SJ, O’Connor WA, Scanes E, Byrne M, et al. Effects of multiple climate change stressors: ocean acidification interacts with warming, hyposalinity, and low food supply on the larvae of the brooding flat oyster *Ostrea angasi*. Mar Biol. 2016;163(5):1–17.

36. Pereira RRC, Scanes E, Parker LM, Byrne M, Cole VJ, Ross PM. Restoring the flat oyster *Ostrea angasi* in the face of a changing climate. Mar Ecol Prog Ser. 2019;625:27–39.

37. Ridgway K. Long-term trend and decadal variability of the southward penetration of the East Australian Current. Geophys Res Lett. 2007;34(13).

38. Hobday AJ, Pecl GT. Identification of global marine hotspots: sentinels for change and vanguards for adaptation action. Rev Fish Biol Fish. 2014;24(2):415–25.

39. Babcock, R.C., Bustamante, R.H., Fulton, E.A., Fulton, D.J., Haywood, M.D., Hobday, A.J., Kenyon, R., Matear, R.J., Plagányi, E.E., Richardson, A.J. and Vanderklift, M.A. Severe continental-scale impacts of climate change are happening now: Extreme climate events impact marine habitat forming communities along 45% of Australia’s coast. Frontiers in Marine Science, 6, 2019;p.411.

40. Schiel, D.R., Steinbeck, J.R. and Foster, M.S.,. Ten years of induced ocean warming causes comprehensive changes in marine benthic communities. Ecology, 85(7), 2004; pp.1833–1839.

41. Nell JA, O’Connor WA. The evaluation of fresh algae and stored algal concentrates as a food source for Sydney rock oyster, *Saccostrea commercialis* (Iredale & Roughley), larvae. Aquaculture. 1991;99(3-4):277–84.

42. Roy P, Williams R, Jones A, Yassini I, Gibbs P, Coates B, et al. Structure and function of south-east Australian estuaries. Estuar Coast Shelf Sci. 2001;53(3):351–84.

43. Lucas A, Beninger PG. The use of physiological condition indices in marine bivalve aquaculture. Aquaculture. 1985;44(3):187–200.

44. Mann R. A comparison of methods for calculating condition index in eastern oysters Crassostrea virginica (Gmelin, 1791). J Shellfish Res. 1992;11(1):55.

45. Bligh EG, Dyer WJ. A rapid method of total lipid extraction and purification. Can J Biochem Physiol. 1959;37(8):911–7.

46. Volkman JK, Nichols PD. Applications of thin layer chromatography-flame ionization detection to the analysis of lipids and pollutants in marine and environmental samples. J Planar Chromatogr. 1991;4:19–26.

47. Ackman R. [11] Flame ionization detection applied to thin-layer chromatography on coated quartz rods. Methods Enzymol. 72: Elsevier; 1981. p. 205–52.

48. Sinanoglou VJ, Strati IF, Bratakos SM, Proestos C, Zoumpoulakis P, Miniadis-Meimaroglou S. On the combined application of Iatroscan TLC-FID and GC-FID to identify total, neutral, and polar lipids and their fatty acids extracted from foods. ISRN Chromatography. 2013;2013.

49. Anderson M, Gorley RN, Clarke RK. Permanova+ for primer: Guide to software and statisticl methods: Primer-E Limited; 2008, Plymouth, UK.

50. Pörtner H-O, Reipschläger A, Heisler N. Acid-base regulation, metabolism and energetics in *Sipunculus nudus* as a function of ambient carbon dioxide level. J Exp Biol. 1998;201(1):43–55.

51. Hiebenthal C, Philipp EE, Eisenhauer A, Wahl M. Effects of seawater *p*CO_2_ and temperature on shell growth, shell stability, condition and cellular stress of Western Baltic Sea *Mytilus edulis* (L.) and *Arctica islandica* (L.). Mar Biol. 2013;160(8):2073–87.

52. Talmage SC, Gobler CJ. Effects of elevated temperature and carbon dioxide on the growth and survival of larvae and juveniles of three species of northwest Atlantic bivalves. PloS one. 2011;6(10):e26941.

53. Bayne B, Bayne C, Carefoot T, Thompson R. The physiological ecology of *Mytilus californianus* Conrad. 1. Metabolism and energy balance. Oecologia. 1976:211–28.

54. Abad, M., Ruiz, C., Martinez, D., Mosquera, G. and Sánchez, J. Seasonal variations of lipid classes and fatty acids in flat oyster, *Ostrea edulis*, from San Cibran (Galicia, Spain). Comparative Biochemistry and Physiology Part C: Pharmacology, Toxicology and Endocrinology, 1995;110(2). pp.109–118.

55. Fields PA, Zuzow MJ, Tomanek L. Proteomic responses of blue mussel (*Mytilus*) congeners to temperature acclimation. J Exp Biol. 2012;215(7):1106–16.

56. Scanes E, Parker LM, O’Connor WA, Gibbs MC, Ross PM. Copper and ocean acidification interact to lower maternal investment, but have little effect on adult physiology of the Sydney rock oyster *Saccostrea glomerata*. Aquat Toxicol. 2018;203:51–60.

57. Somero GN. Thermal physiology and vertical zonation of intertidal animals: optima, limits, and costs of living. Integr Comp Biol. 2002;42(4):780–9.

58. Anestis A, Lazou A, Pörtner HO, Michaelidis B. Behavioral, metabolic, and molecular stress responses of marine bivalve *Mytilus galloprovincialis* during long-term acclimation at increasing ambient temperature. American Journal of Physiology-Regulatory, Integr Comp Physiol. 2007;293(2):R911–R21.

59. Sørensen JG, Kristensen TN, Loeschcke V. The evolutionary and ecological role of heat shock proteins. Ecol Lett. 2003;6(11):1025–37.

60. Ivanina A, Taylor C, Sokolova I. Effects of elevated temperature and cadmium exposure on stress protein response in eastern oysters *Crassostrea virginica* (Gmelin). Aquat Toxicol. 2009;91(3):245–54.

61. Abele D, Heise K, Pörtner H-O, Puntarulo S. Temperature-dependence of mitochondrial function and production of reactive oxygen species in the intertidal mud clam *Mya arenaria*. J Exp Biol. 2002;205(13):1831–41.

62. Somero, G.N. The physiology of global change: linking patterns to mechanisms. Annual Review of Marine Science, 2012;4, pp.39–61.

63. Rivest, E.B., Comeau, S. and Cornwall, C.E. The role of natural variability in shaping the response of coral reef organisms to climate change. Current Climate Change Reports, 2017;3(4), pp.271–281.

64. McAfee, D., O’connor, W.A. and Bishop, M.J. Fast-growing oysters show reduced capacity to provide a thermal refuge to intertidal biodiversity at high temperatures. Journal of Animal Ecology, 2017;86(6), pp.1352–1362.

65. Mitchell, I.M., Crawford, C.M. and Rushton, M.J. Flat oyster (Ostrea angasi) growth and survival rates at Georges Bay, Tasmania (Australia). Aquaculture, 2000;191(4), pp.309–321.

66. Lipcius RN, Burke RP. Successful recruitment, survival and long-term persistence of eastern oyster and hooked mussel on a subtidal, artificial restoration reef system in Chesapeake Bay. PloS one. 2018;13(10):e0204329.

67. Laing I, Walker P, Areal F. Return of the native–is European oyster (*Ostrea edulis*) stock restoration in the UK feasible? Aquatic Living Resources. 2006;19(3):283–7.

68. Rodolfo-Metalpa R, Lombardi C, Cocito S, Hall-Spencer JM, Gambi MC. Effects of ocean acidification and high temperatures on the bryozoan *Myriapora truncata* at natural CO_2_ vents. Mar Ecol. 2010;31(3):447–56.

69. Calosi P, Rastrick S, Graziano M, Thomas S, Baggini C, Carter H, et al. Distribution of sea urchins living near shallow water CO_2_ vents is dependent upon species acid–base and ion-regulatory abilities. Mar Pollut Bull. 2013;73(2):470–84.

